# Immunoblot-based activity assay for heme-containing histidine kinases

**DOI:** 10.1101/2025.05.28.656737

**Authors:** Grant Larson, Ambika Bhagi-Damodaran, Anoop Rama Damodaran

## Abstract

Histidine kinases (HKs) are essential bacterial signal transduction proteins and attractive drug targets due to their critical cellular functions. Direct measurement of their autophosphorylation activity is crucial for developing inhibitors and advancing disease treatments. While [*γ*-^32^P]-ATP radiolabeling has long been a conventional method for kinase activity measurements, its reliance on ^32^P introduces inherent limitations. The isotope’s short half-life imposes time-sensitive constraints on experiments, and stringent radiation safety and compliance requirements significantly increase operational costs and hinder scalability. Several alternative methods, including fluorescent phosphate-binding probes and fluorescence- or luminescence-based antibodies, have been developed for HKs that overcome these challenges. However, for heme-based HKs, these alternatives suffer from interference due to intrinsic heme fluorescence and luminescence. In this work, we address this interference by employing a near-IR fluorophore-labeled secondary antibody. Combined with an ATP*γ*S immunoblot-based method, we demonstrate semi-quantitative detection of thiophosphorylation activities of *Mycobacterium tuberculosis* DosS (a prototypical heme-based HK) in different ligation states. Consistent with previous radiolabeling studies, we show DosS’s ability for auto-thiophosphorylation is significantly enhanced with CO as compared to O_2_. This low-cost, easy-to-implement method simplifies and democratizes heme-based HK activity measurement.

## Introduction

Histidine kinases (HKs) are ubiquitous signal transduction proteins found in bacteria, fungi, and plants, where they play a crucial role in environmental sensing and adaptive gene regulation.^1,2^ These proteins typically consist of a sensor domain, a linker domain, a dimerization and histidine phosphotransfer (DHp) domain, and a catalytic/ATP-binding (CA) domain. Upon detecting a stimulus, the sensor domain triggers conformational changes that propagate through the linker and DHp domains, leading to auto-phosphorylation at a conserved histidine residue within the DHp domain.^3^ The phosphoryl group is then transferred to a response regulator, which modulates gene transcription by directly binding to the organism’s DNA^4^ (**Fig. S1**). Given the essential role of HKs in bacterial signaling and adaptation, they represent promising therapeutic targets for antibiotic discovery.^5^ Developing HK inhibitors requires reliable, accessible, and scalable autophosphorylation assays. Radiometric assays that employ [*γ*-^32^P]-ATP have been generally considered as the gold standard for kinase activity (**Fig. S2**).^6,7^ However, the short half-life of ^32^P coupled with radiation safety and compliance challenges along with high reagent and operational costs have motivated the development of alternative approaches. In turn, approaches including fluorescence-or luminescence-based phosphohistidine antibodies,^8^ and fluorescent^9^ or electrophoretic mobility^10–12^-based phosphate-binding probes have been developed.

Despite the development of various alternative approaches, heme-based HKs have predominantly been investigated using the radiometric approach.^13,14^ A select few studies have employed electrophoretic mobility-based assays that utilize gels loaded with phosphate-binding molecules.^11,12^ Fluorescence-or luminescence-based kinase assays have not been employed for heme-based HKs, likely due to interference caused by the strong intrinsic fluorescence and luminescence of the heme-cofactor. Even a minor fraction of retained heme that persists after SDS denaturation of heme-based HK and gel migration can introduce significant intrinsic fluorescence interference under standard visible-light fluorescent detection settings. Furthermore, the low stability of phosphorylated histidine^15^ precludes the use of thermal denaturation or acidification protocols for complete heme removal. In this work, we demonstrate that fluorescent detection of kinase activity in heme-based HKs is achievable by employing a near-IR (NIR) fluorophore-labeled secondary antibody, which circumvents interference from the heme cofactor’s intrinsic fluorescence and luminescence. Combining NIR fluorescent detection with a previously developed ATP*γ*S-based immunoblotting method^16,17^ for detecting thiophosphohistidine (tpHis, a phosphohistidine analog with significantly enhanced stability^15^), we provide a robust fluorescent-based kinase activity assay that is reliable, accessible and scalable. We apply this method to demonstrate semi-quantitative detection of thiophosphorylation activities for *Mycobacterium tuberculosis* DosS, a prototypical heme-based HK whose activity depends on the ligation state of its heme sensor.^14^ Consistent with previous radiolabeling assays, our studies show that DosS’s auto-thiophosphorylation activity is significantly enhanced when its heme sensor is bound to CO as compared to O_2_.

## Results and Discussion

To demonstrate the unsuitability of kinase assays that employ standard visible-light fluorescent or luminescent detection for heme-based HKs, we ran an SDS PAGE gel with a set of samples that included unreacted ferric and ferrous DosS, and CO/O_2_-bound ferrous DosS reacted with ATP for up to 60 minutes (**Fig. S3**). We also included a heme-free HK construct ^_^ a truncated form of *T. maritima* HK853^18^ that lacks its sensor domain and undergoes auto-phosphorylation even in the absence of a signal. Upon imaging the gel using visible-light fluorescent detection parameters prior to any fluorescent staining (**Fig. 1a**), we observe a band corresponding to a strong and uniform fluorescence signal across all DosS samples (MW=64 kDa). On the other hand, HK853 which possesses no intrinsic fluorescence under these imaging conditions, reveal clear lanes and the absence of any band at positions corresponding to its molecular weight of 32 kDa (**Fig. S4**). Upon staining the gel with PhosTag Aqua stain^10^ (a phosphate-binding fluorophore) to assess autophosphorylation levels, we observe that intrinsic fluorescence obscures DosS signals (**Fig. 1 b**, upper arrow). In contrast, HK853 reveals increasing auto-phosphorylation over the time course of the reaction (**Fig. 1b**, lower arrow). Next, we ran a duplicate SDS PAGE gel, transferred it to a nitrocellulose membrane and imaged the membrane with luminol using standard chemiluminescent detection without having carried out any immunoblotting with primary/secondary antibodies. We note that the DosS band is able to catalyze luminol oxidation generating strong and unwanted luminescence (**Fig. 1c**, upper arrow). Again, HK853 reveals clear lanes and the absence of any luminescent bands at positions corresponding to its molecular weight (**Fig. 1c**, lower arrow). Taken together, these studies show that even a minor fraction of retained heme that persists after SDS denaturation of heme-based HKs and gel migration can render standard visible-light fluorescent and luminescent detection-based kinase assays unsuitable for heme-based HKs.

**Figure 1.**
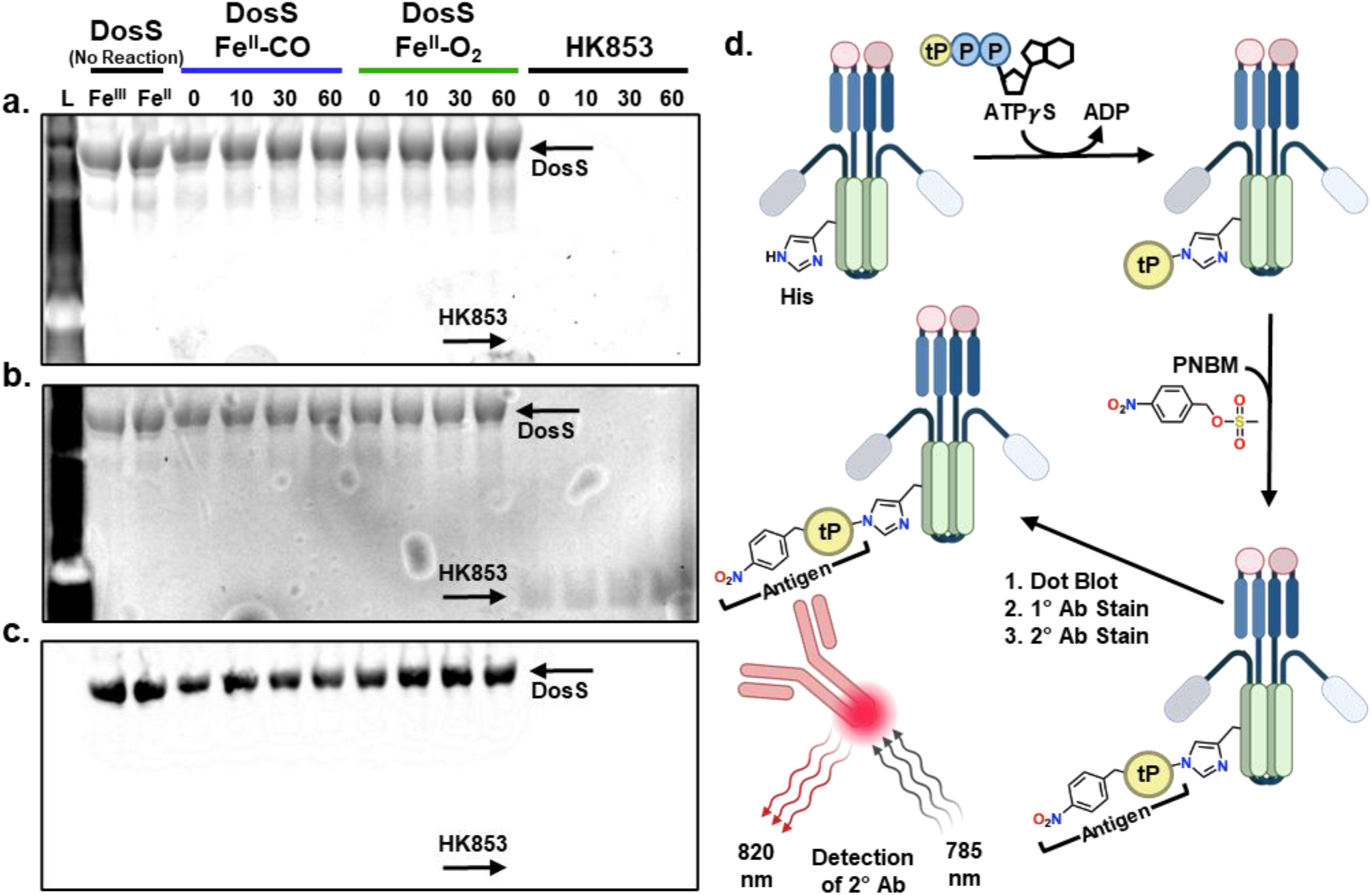
(a) Fluorescent image of SDS PAGE gel demonstrating unreacted ferric and ferrous DosS along with ferrous-CO, ferrous-O_2_, and HK853 reacted with ATP upto 60 minutes. L indicates the lane for protein MW ladder. The fluorescence emission from the gel was measured with excitation at 532 nm and emission at 553 nm right after electrophoretic separation. (b) Gel in (a) was stained with PhosTag Aqua and imaged again with excitation at 532 nm and emission at 553 nm. (c) Luminescent image of SDS PAGE gel that is duplicate of that in (a) but also subjected to western blot transfer and reacted with luminol. (d) Schematic illustrating the NIR fluorescence-based thiophosphorylation developed in this work. The HK reacts with ATP*γ*S, which transfers its thiophosphate (tP) group to the histidine residue of HK. The tP is chemically modified with PNBM to form the PNB-tP antigen. This modified protein sample is dot blotted onto a nitrocellulose membrane and stained with primary and secondary antibodies. Our method makes use of a secondary antibody conjugated to a NIR labelled fluorophore, allowing for detection without interference from the heme. Only one antibody is shown for clarity.

Given that heme does not absorb in the NIR, kinase assays employing secondary antibodies conjugated to NIR fluorophores can circumvent challenges related to the intrinsic fluorescence from residual heme in denatured DosS samples. We chose an immunoblot method (**Fig. 1d, Fig. S2**) that uses ATP*γ*S instead of ATP and detects tpHis by converting it to the biorthogonal epitope para-nitrobenzyl thiophosphoester (PNB-tP), which then binds strongly to an anti-thiophosphate ester primary antibody.^16,17^ This method has been utilized to detect and probe numerous kinases,^16,17^ and offers several advantages over using phosphorylated-histidine detecting antibodies. First, the method is universal and applicable to histidine, serine, threonine, and tyrosine kinases. Second, focusing on HKs, tpHis is much more stable than phosphohistidine,^19,20^ thereby mitigating potential hydrolysis related challenges during subsequent immunoblotting steps that could otherwise lead to unreliable activity measurements. We began by investigating the time-dependent thiophosphorylation of DosS in its CO- and O_2_-bound forms. We included HK853 as positive control and a useful benchmark for assessing the relative activity of DosS under different conditions. The thiophosphorylation reactions were quenched with EDTA at predetermined time points and were subsequently reacted with p-nitro benzyl mesylate (PNBM). The protein samples were then subjected to SDS-PAGE for electrophoretic separation and western blot transfer (**Fig. 2a-b**). As the samples were not boiled during denaturation with gel loading buffer to prevent any hydrolysis of thiophosphohistidine, two distinct bands appeared in the DosS lanes corresponding to the dimeric (upper band, 128 kDa) and monomeric (lower band, 64 kDa) species. The band intensities of the DosS dimeric and monomeric species increased over time, indicative of auto-thiophosphorylation activity. Signals from both bands were summed to obtain the total auto-thiophosphorylation for each time point and then normalized to the auto-thiophosphorylation signal observed for HK853 at the 30-minute time point. These measurements reveal that DosS Fe^II^-CO exhibits an auto-thiophosphorylation rate approximately six times greater than that of DosS Fe^II^-O_2_, a trend consistent with previous^13,21^ radiolabeling-based studies.

**Figure 2.**
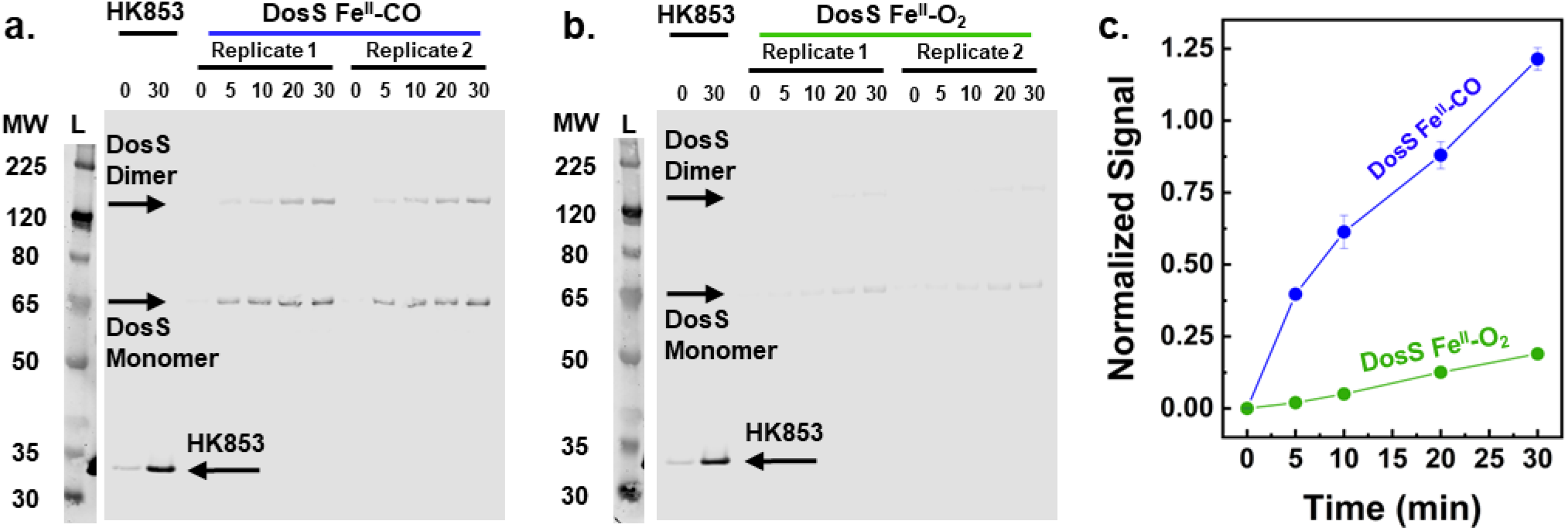
Fluorescent image of a western blot containing (a) DosS Fe^II^-CO and (b) DosS Fe^II^-O_2_ thiophosphorylated samples collected over 0 to 30 minutes, detected using a NIR fluorophore-labelled secondary antibody. L indicates the lane for protein MW ladder. a-b also contain lanes corresponding to HK853’s reaction with ATPγS quenched at 0 and 30 min. The fluorescence emission from the membrane was measured with excitation at 785 nm and emission at 820 nm (c) Signal from thiophosphorylation of CO-(blue) and O_2_-bound (green) DosS in a-b plotted with respect to time. The signal is normalized to HK853. Blue and green lines serve as guide to the eye.

Having overcome challenges related to intrinsic fluorescence of heme by NIR fluorescent immunoblotting, we proceeded to do additional auto-thiophosphorylation measurements in a dot blot format, a technique that bypasses SDS-PAGE by directly applying purified protein samples to a membrane. Traditional dot blotting methods, however, can produce irregularly shaped or unevenly sized dots, making quantification difficult. To improve consistency and precision, we utilized a 96-well microfiltration Bio-Dot apparatus, which ensures uniform spot size and spacing across the membrane. This approach offers several advantages including significantly increased throughput as up to 96 samples can be analyzed on a single membrane. More importantly, the use of uniformly sized dots facilitates reproducibility, making dot blotting a practical and scalable route for HK activity assays. We conducted time-dependent thiophosphorylation of DosS in its CO- and O_2_-bound forms along with positive control HK853 for kinase activity investigations in the dot-blot format. All samples were applied to the same membrane to ensure consistency, allowing for uniform staining with a primary antibody specific to PNB-tpHis and a NIR fluorophore-labeled secondary antibody. We observe that the fluorescence intensity of both CO and O_2_-bound DosS samples progressively increase from left to right, corresponding to time-dependent increase in auto-thiophosphorylation signal (**Fig. 3a, Fig. S5**). Overall, these measurements also reveal that DosS Fe^II^-CO (**Fig. 3b** blue circles) exhibits an enhanced auto-thiophosphorylation rate approximately five times greater than DosS Fe^II^-O_2_ (**Fig. 3b** green circles). Next, we performed time-dependent authothiophosporylation studies with CO and O_2_-bound forms of H395Q DosS (an inactive DosS mutant). This mutant retains the heme-binding domain but lacks the target H395 histidine residue that undergoes thiophosphorylation in WT DosS and serves as a negative control. As expected, we observe no detectable thiophosphorylation in H395Q DosS across both ligation states (open blue circles for Fe^II^-CO and open green circles for Fe^II^-O_2_ forms in **Fig. 3b, Fig. S6**). This confirms that the signals measured in our auto-thiophosphorylation assays are due to functional thiophosphorylation events at the H395 position. Overall, these studies reveal the potential of our NIR fluorescence-based thiophosphorylation measurements to characterize the kinase activity of heme-based HKs.

**Figure 3.**
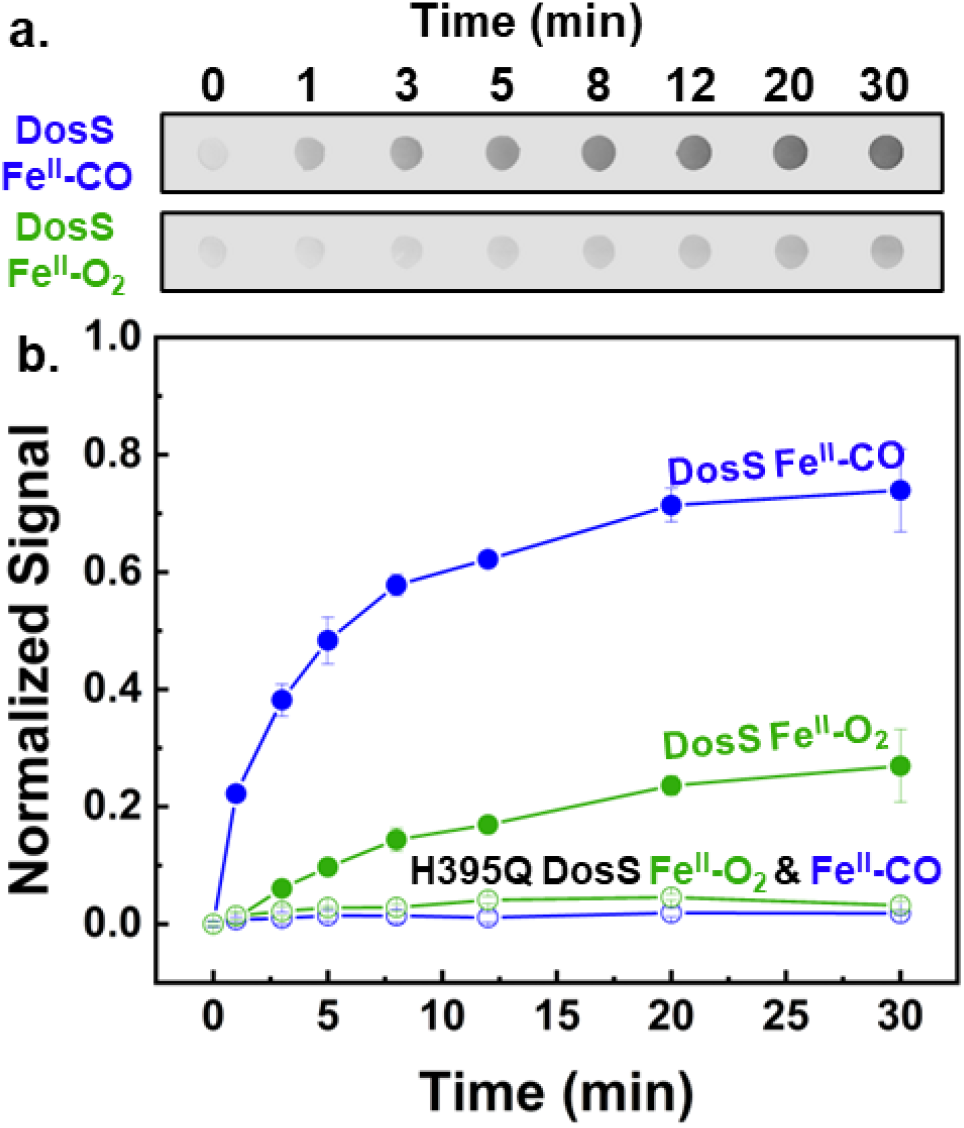
(a) Fluorescent images of portions of an immunostained nitrocellulose membrane containing active DosS Fe^II^-CO (darker spots on top) and DosS Fe^II^-O_2_ (lighter spots on bottom). The fluorescence emission was measured with excitation at 785 nm and emission at 820 nm (b) Signal from thiophosphorylation of CO-(blue circles) and O_2_-bound (green circles) DosS plotted with respect to time. Open circles represent signals from CO-(blue circles) and O_2_-bound (green circles) H395Q DosS plotted with respect to time. The signals are normalized to HK853. Blue and green lines serve as guide to the eye.

## Conclusion

Microbes utilize a diverse array of heme-based kinases to sense and signal changes in their redox environment.^4^ Understanding how these kinases differentiate between various redox stimuli and efficiently transmit signals downstream to response regulators is a critical research focus from both biochemical and microbiological perspectives. Furthermore, the involvement of DosS-like HKs in infectious diseases makes them appealing drug targets.^22^ This work introduces a NIR fluorescence-based thiophosphorylation assay to investigate the kinase activity of heme-based HKs in a low-cost, easy-to-implement, and scalable format. We believe that this assay paves the way for more rigorous characterization of heme-based HKs as well as the development of high-throughput modalities to discover their small-molecule inhibitors.

## Acknowledgement

GL, ABD, and ARD acknowledge the support of NIH NIGMS grant #R35GM138277 for funding this work

## Author Contributions

ARD and ABD designed and supervised this study. GL developed and performed all the biochemical assays. GL, ABD, and ARD wrote the manuscript with contributions from all authors. All authors have given approval to the final version of the manuscript.

## Methods

### DosS Expression

BL21(DE3) *E. coli* cells (ThermoFisher Scientific) were transformed with a pET23a(+) plasmid containing a gene for DosS (Accession: P9WGK3) with an N-terminal 6xHis tag and a TEV protease site (GenScript) and a pACYC-GroEL/ES-TF plasmid containing the chaperonin GroEL/ES (Addgene 83923). Transformed cells were plated on LB agar plates containing 100 μg/mL ampicillin and 37 μg/mL chloramphenicol. Primary cultures (100 mL 2XYT media containing 100 μg/mL ampicillin and 37 μg/mL chloramphenicol) were inoculated with a single colony from an agar plate. They were grown overnight at 37 °C while shaking at 220 rpm. Secondary cultures (1 L 2XYT media containing 100 μg/mL ampicillin and 37 μg/mL chloramphenicol) were inoculated with 25 mL of primary culture and were grown at 37 °C while shaking at 220 rpm. The cell density was monitored until the OD_600_ reached 0.5 – 0.7. Afterwards, the shaker conditions were changed to 30 °C and 170 rpm. Heme precursor molecules, 5-aminolevulinic acid and ferrous ammonium sulfate, were added to the cultures at 0.5 mM and 0.1 mM, respectively. Protein expression was induced with 0.5 mM isopropyl-1-thio-β-D-galactopyranoside (IPTG, GoldBio) and was allowed to proceed for 18 hours. Cells were harvested via centrifugation at 2000 rpm for 30 min at 4 °C (Beckman Coulter).

### HK853 Expression

BL21(DE3) *E. coli* cells were transformed with a pET28a(+) plasmid (generously provided by the Carlson lab at the University of Minnesota) containing a gene for the cytosolic portion of HK853 (Accession: Q9WZV7; amino acids 234 – 489) with an N-terminal 6xHis tag and a thrombin protease site (GenScript). Transformed cells were plated on LB agar plates containing 50 μg/mL kanamycin. Primary cultures (100 mL 2XYT media containing 50 μg/mL kanamycin) were inoculated with a single colony from an agar plate. They were grown overnight at 37 °C while shaking at 220 rpm. Secondary cultures (1 L 2XYT media containing 50 μg/mL kanamycin) were inoculated with 25 mL of primary culture and were grown at 37 °C while shaking at 220 rpm. The cell density was monitored until the OD_600_ reached 0.5 – 0.7. Afterwards, the shaker conditions were changed to 18 °C and 170 rpm. Protein expression was induced with 0.5 mM IPTG and was allowed to proceed for 18 hours. Cells were harvested via centrifugation at 2000 rpm for 30 min at 4 °C (Beckman Coulter).

### DosS and HK853 Purification

Cell pellets were resuspended in 50 mM Tris (pH 7.5), 250 mM sodium chloride, and 1% Triton X-100 and lysed via sonification (Fisher Scientific) while on ice. The crude lysate was centrifuged for 1 hour at 20,000 rpm and 4 °C (Beckman Coulter). The supernatant was filtered using a 0.44 μm syringe filter (Sartorius) and applied to a 5 mL HisTrap FF column (GE Healthcare) at a rate of 2 mL/min. The column was washed with 75 mL 50 mM Tris (pH 7.5), 250 mM sodium chloride, and 20 mM imidazole at a rate of 2 mL/min. Recombinant protein was eluted with 50 mM Tris (pH 7.5), 250 mM sodium chloride, and 400 mM imidazole. The semi-pure DosS solution was oxidized with 2.5 mM potassium ferricyanide (III). Excess oxidant was removed by buffer exchanging the protein solution into 50 mM Tris pH 8.0, 100 mM NaCl, and 5% glycerol via gel filtration through a PD-10 column (Cytiva). A concentrated protein solution was purified via size exclusion chromatography (SEC) with a HiPrep 16/60 Sephacryl S-200 HR column (GE Healthcare). The final DosS concentration was determined via UV-Vis absorbance at 406 nm using the extinction coefficient 145 mM^-1^ cm^-1^ determined via hemochromogen assay.^23^ Aliquots of the protein solution were flash-frozen in liquid nitrogen and stored at -80 °C.

A concentrated solution of HK853 was purified via SEC with a HiLoad 26/600 Superdex 75 pg column (GE Healthcare). The final HK853 concentration was determined via UV-Vis absorbance at 280 nm using the EXPASY predicted extinction coefficient 21430 M^-1^ cm^-1^. Aliquots of the protein solution were flash-frozen in liquid nitrogen and stored at -80 °C.

### Autophosphorylation Assay

Solutions of Ferrous DosS were prepared anaerobically (Coy Lab Products) by reducing Ferric DosS with sodium dithionite. Ferrous-CO DosS was prepared by adding 20 molar equivalents of CORM-A1 (Sigma Aldrich). Ferrous-O_2_ DosS was prepared by exposing Ferrous DosS to air for 5 minutes. All redox and ligation states of DosS were confirmed via UV-Vis (Agilent). Autophosphorylation reactions were performed in 50 mM Tris (pH 8.0), 50 mM potassium chloride, 10 mM magnesium (II) chloride, and 10% glycerol while shaking at 300 rpm at 20 °C. Reactions contained 5 uM HK853 or DosS and were initiated by adding 1 mM adenosine 5′-[γ-thio]triphosphate (AGS). At designated time points, aliquots of the reaction mixture were removed and quenched by adding 100 mM EDTA. The autophosphorylated proteins in the quenched reaction mixtures were modified with 3 mM para-nitrobenzyl mesylate (PNBM, Astatech). The modification reaction proceeded for 90 minutes, shaking at 300 rpm at 20 °C.

### Dot Blot of Autophosphorylation Samples

A nitrocellulose membrane was wetted with 20 mM Tris (pH 8.0) and 150 mM sodium chloride (TBS). The membrane was loaded into a 96-well Bio-Dot apparatus (Bio-Rad), which was set up following the manufacturer’s instructions. PNBM-modified protein solutions were diluted to concentrations of 0.1 μM, and 200 μL were loaded into each well. After gravity filtration of the protein solution through the nitrocellulose membrane, the membrane was removed from the apparatus and dried at 37 °C for 30 min to fix the proteins to the membrane.

### Western Blot of Autophosphorylation Samples

Proteins were separated via polyacrylamide gel electrophoresis using an 8% Tris-Glycine gel and Tris-Glycine running buffer (pH 8.8). The proteins were transferred to a nitrocellulose membrane, which was dried at 37 °C for 30 min to fix the proteins to the membrane.

### Immunoblotting Autophosphorylation Samples

Membranes from Dot Blotting and Western Blotting were rehydrated with TBS and blocked with Intercept (TBS) protein-free blocking buffer (LI-COR) for 60 min at room temperature while rocking. Primary antibody solution was prepared by diluting rabbit anti-thiophosphoester antibody (Abcam, ab92570) in Intercept T20 (TBS) protein-free antibody diluent. The membranes were incubated with primary antibody solution for 16 hours at 4 °C while rocking, washed four times with TBST (TBS + 0.1% Tween 20), and incubated with the secondary antibody solution containing goat anti-rabbit IgG Alexa Fluor Plus 800 (ThermoFisher Scientific, A32735), for 60 minutes at room temperature. The membranes were washed four times with TBST and dried at 37 °C for 60 minutes before imaging on an Odyssey M imager (LI-COR) using the 800 channel. The images were processed and analyzed using Empiria Studio 3.0 (LI-COR). Subsequently, the membranes were stained with SYPRO Ruby protein blot stain (ThermoFisher Scientific) to determine the amount of total protein. The stained membranes were then imaged with an Odyssey M imager using the 488A channel. The images were processed and analyzed using Empiria Studio 3.0.

