## Supplementary material for "Immunoblot-based activity assay for heme-containing histidine kinases": SI

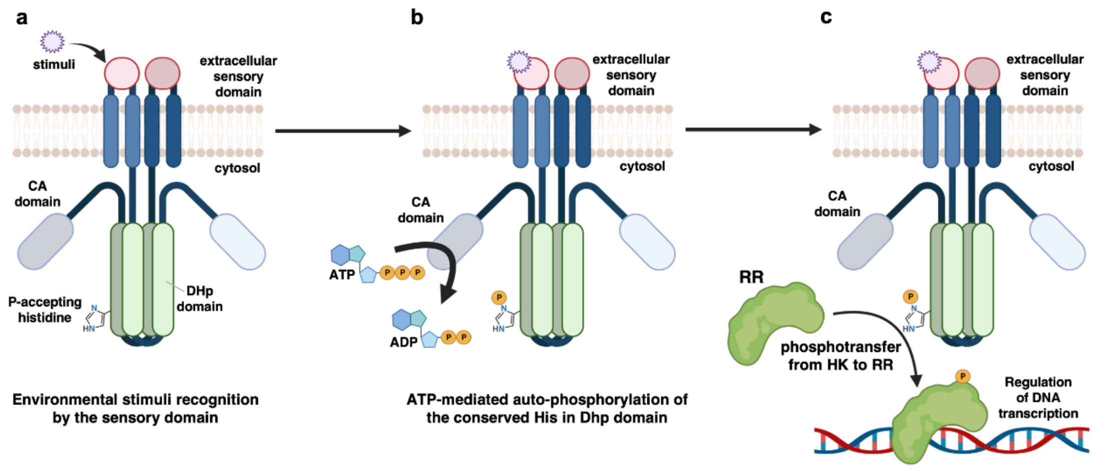


**Supplemental Figure 1.** A simplistic model of a Histidine Kinase (HK) and Response Regulator (RR) -mediated bacterial TCS pathway. (a) An environmental stimulus is recognized by the HK's extracellular sensory domain. (b) Stimulus recognition by the sensory domain initiates ATP-mediated auto-phosphorylation of a conserved histidine in DHp domain. (c) The signal is further transmitted via phosphotransfer from HK to RR. The phosphorylated RR binds to the DNA and alters gene transcription to allow for cellular adaptation to the environmental stimuli. Note: This model demonstrates a cis auto-phosphorylation mechanism in panel b) i.e. CA domain autophosphorylates DHp on the same HK monomer. The mechanism of DosS autophosphorylation is not known.

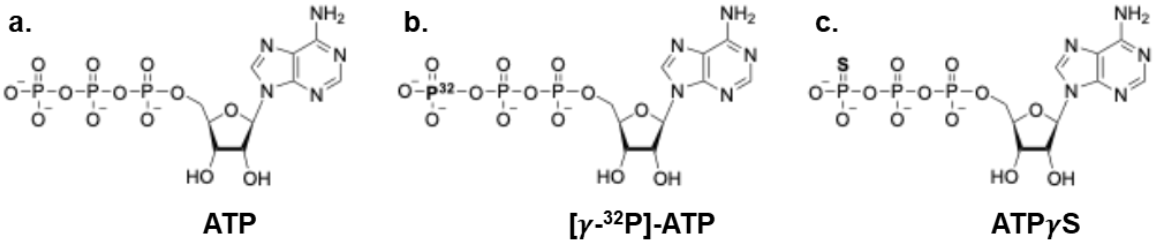


**Supplemental Figure 2.** Structures of (a) ATP, (b) [$\gamma$-$^{32}$P]-ATP, and (c) ATP$\gamma$S. Structural differences in (b) and (c) are bolded.

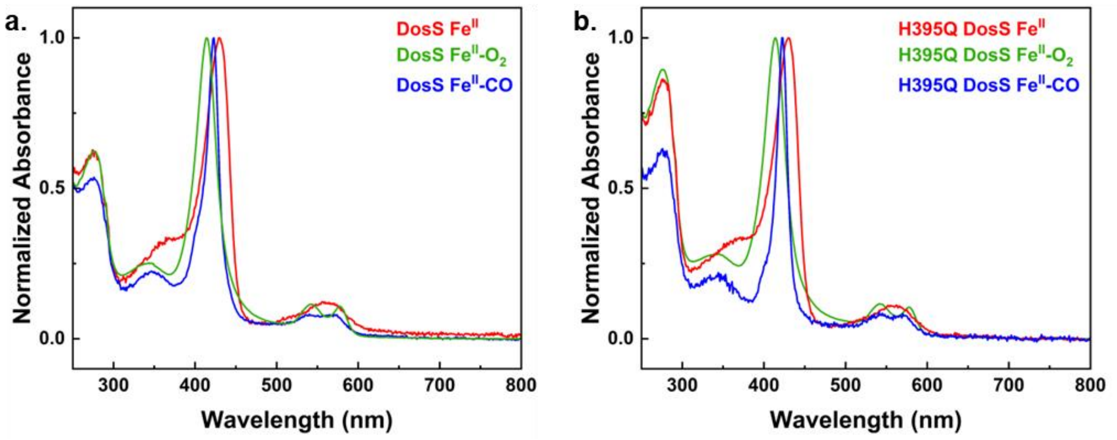


**Supplemental Figure 3.** UV-Vis spectra of (a) wild-type DosS and (b) H395Q DosS with various ligands. Soret maxima of DosS samples are as follows: DosS $Fe^{II}$-$O_2$ (414 nm), DosS $Fe^{II}$-CO (422 nm), DosS $Fe^{II}$ (430 nm).

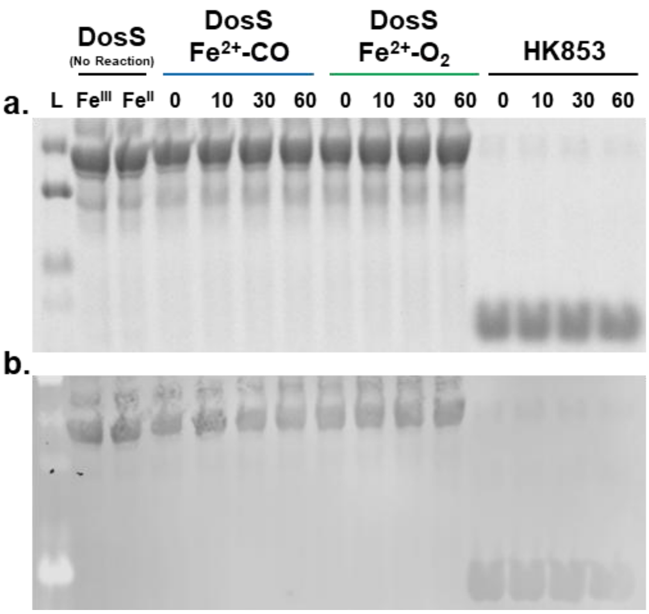


**Supplemental Figure 4.** (a) The coomassie-stained gel shown in Figure 1a and 1b to demonstrate the presence of HK853 and DosS in equivalent amounts. (b) The SYPRO Ruby stained membrane shown in Figure 1c to demonstrate the presence of HK853 and DosS in equivalent amounts.

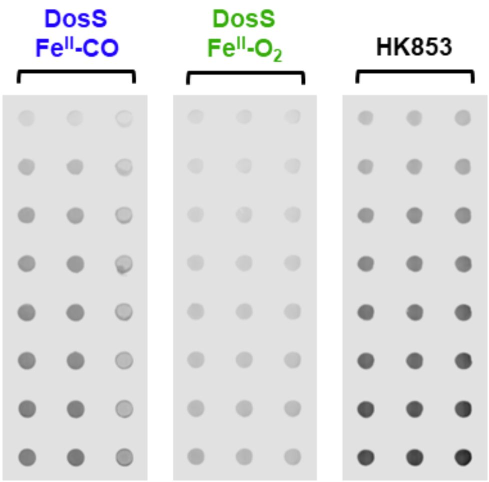


**Supplemental Figure 5.** Fluorescent image of nitrocellulose membrane blotted with CO and $O_2$-bound DosS and HK853 samples. The third column of DosS $Fe^{II}$-CO samples were excluded from data analysis due to irregularities in the dots. The fluorescence emission was measured with excitation at 785 nm and emission at 820 nm.

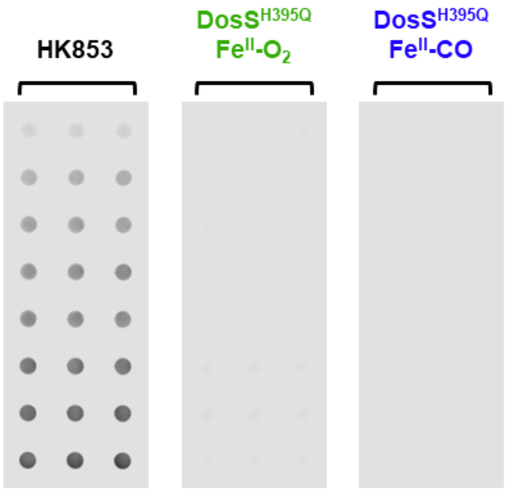


**Supplemental Figure 6.** Fluorescent image of nitrocellulose membrane blotted with HK853, and $O_2$, CO-bound H395Q DosS samples. The fluorescence emission was measured with excitation at 785 nm and emission at 820 nm
